# Voxel-Wise Brain Graphs from Diffusion MRI: Intrinsic Eigenspace Dimensionality and Application to Functional MRI

**DOI:** 10.1101/2022.09.29.510097

**Authors:** Hamid Behjat, Anjali Tarun, David Abramian, Martin Larsson, Dimitri Van De Ville

**Author notes:** H. Behjat and A. Tarun provided equal contribution.

## Abstract

Structural brain graphs are conventionally limited to defining nodes as gray matter regions from an atlas, with edges reflecting the density of axonal projections between pairs of nodes. Here we explicitly model the entire set of voxels within a brain mask as nodes of high-resolution, subject-specific graphs. We define the strength of local voxel-to-voxel connections using diffusion tensors and orientation distribution functions derived from diffusion MRI data. We study the graphs’ Laplacian spectral properties on data from the Human Connectome Project. We then assess the extent of inter-subject variability of the Laplacian eigenmodes via a procrustes validation scheme. Finally, we demonstrate the extent to which functional MRI data are shaped by the underlying anatomical structure via graph signal processing. The graph Laplacian eigen-modes manifest highly resolved spatial profiles, reflecting distributed patterns that correspond to major white matter pathways. We show that the intrinsic dimensionality of the eigenspace of such high-resolution graphs is only a mere fraction of the graph dimensions. By projecting task and resting-state data on low-frequency graph Laplacian eigenmodes, we show that brain activity can be well approximated by a small subset of low-frequency components. The proposed graphs open new avenues in studying the brain, be it, by exploring their organisational properties via graph or spectral graph theory, or by treating them as the scaffold on which brain function is observed at the individual level.

## 1. Introduction

Magnetic resonance imaging (MRI) has provided an effective means to map the brain’s anatomical scaffold, using diffusion MRI, and, in parallel, to track brain neural activity using functional MRI (fMRI). Extensive datasets that include diffusion and functional data on the same set of subjects, such as the Human Connectome Project (HCP) (Essen et al., 2013), have been made freely available, and with such accessibility, various methodological developments that aim to integrate the two modalities have emerged.

Computational neuroimaging has successfully adopted graph theory, creating a new field of interdisciplinary research called *network neuroscience* (Bassett and Sporns, 2017). Structural and functional *connectomes* are independently defined, and consequently analyzed using graph theory measures to provide a better understanding of their organizing network principles (Bullmore and Sporns, 2009). There is also an increasing interest in deciphering how underlying brain anatomy supports the emergence of spatially and temporally varying distributed patterns of functional activity (Mišić et al., 2015; Iraji et al., 2019; Suárez et al., 2020). In this perspective, it is fitting to consider advancing network neuroscience to a more unifying analysis approach that accounts for the interplay between brain structure and function. The study of signal propagation on structural connectomes (Vézquez-Rodríguez et al., 2020; Weninger et al., 2022) is an example avenue of research that is gaining momentum. Leveraging principles from the recently emerged field of graph signal processing (GSP) (Shuman et al., 2013; Ortega et al., 2018), an alternative framework is taking form, in which functional data are interpreted as functions defined atop of a graph that describes the morphological or wiring structure of the brain, and, in turn, processed using spectral methods that are informed by the underlying brain structure.

GSP generalizes principles from classical discrete signal processing for time series to data defined on irregular domains. GSP has found numerous applications across multiple domains—see e.g. (Ortega et al., 2018) for a recent review, and in particular within neuroimaging, examples include: brain state decoding (Petrantonakis and Kompatsiaris, 2018; Ghoroghchian et al., 2020; Cattai et al., 2021), brain signal denoising (Einizade and Sardouie, 2022), brain activation mapping (Behjat et al., 2015; Abramian et al., 2021; Behjat et al., 2021), source localization (Hyde et al., 2019), diagnosing neuropathology (Itani and Thanou, 2021), tracking fast spatiotemporal cortical dynamics (Glomb et al., 2020; Rué-Queralt et al., 2021), brain fingerprinting and task decoding (Griffa et al., 2022) via quantifying the degree of coupling between brain function and structure (Preti and Van De Ville, 2019), identifying dynamically evolving populations of neurons (Charles et al., 2022), deciphering signatures of attention switching (Huang et al., 2018), manifesting white matter pathways that mediate cortical activity (Tarun et al., 2020), and elucidating perturbations of consciousness induced by brain injury or drugs (Atasoy et al., 2017; Luppi et al., 2022).

The majority of existing applications of GSP in functional brain imaging are limited to region-wise analyses, where regions are defined using a priori brain atlases having 100 to 1000 regions. Motivated by the promising results from existing region-wise GSP studies in linking brain structure and function, and novel recent methods for high-resolution connectomics (Mansour-L et al., 2021) and activation mapping (Zhao et al., 2022), it is fitting to further explore the benefits of large-scale brain graphs at the resolution of *voxels*. Here we present models to derive the strength of connection between adjacent voxels using diffusion MRI data, one based on tensors estimated from diffusion tensor imaging (DTI) (Basser et al., 1994), and the other based on diffusion orientation distribution functions (ODF) estimated from high angular resolution diffusion imaging (HARDI) data (Tuch, 2004); we consider the two signal representation settings to make the methodology applicable to a larger set of available diffusion MRI data. Using data of 100 subjects from the HCP (Essen et al., 2013), we validate the graphs via studying their nodal and spectral measures. We then probe the intrinsic dimensionality of their eigenspace using a Procrustes validation scheme that characterizes inter-subject variability. Finally, we demonstrate the relevance of such high-spatial-resolution voxel-wise graphs within a GSP setting, particularly through studying the energy spectral density of resting-state and task fMRI data on these graphs. We conclude the paper by discussing potential future research avenues in using voxel-wise graphs, in particular, in studying the interaction between brain structure and function.

## 2. Materials and Methods

### 2.A. Datasets

We used MRI data from the publicly available HCP dataset—the 100 unrelated subjects, WU-Minn Consortium (Essen et al., 2013). MRI acquisition protocols of the dataset and preprocessing guidelines for diffusion MRI are extensively described elsewhere (Glasser et al., 2013). We used the minimally preprocessed diffusion and anatomical data. The resting-state and task fMRI scans of each subject were realigned to their mean images, and were registered and resampled onto the diffusion data through rigid-body registration using SPM^1^. Two signal reconstruction methods were applied to the diffusion data: (1) DTI tensor fitting using FSL^2^, and (2) ODF estimations using DSI Studio^3^.

### 2.B. Graphs and their spectra

Let 𝒢 = (𝒩, **A**) denote an undirected, single-connected, weighted graph, consisting of a node set 𝒩, where | 𝒩|= *N*, and a symmetric *N × N* weighted adjacency matrix **A**, wherein any of its nonzero elements *a*_*ij*_ represent the weight of an edge (*i, j*) in the graph. The normalized graph Laplacian (Chung, 1997) is defined as

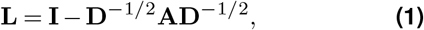

where **I** denotes the identity matrix, and **D** denotes the graph degree matrix, which is diagonal with elements 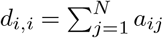. **L** can be diagonalized as **L = U∧U**^**T**^, where **U** = [**u**_1_, **u**_2_, …, **u**_*N*_] is an orthonormal matrix stacking the eigenvectors **u**_*i*_, *i* = 1, …, *N*, and Λ is a diagonal matrix stacking the corresponding eigenvalues Λ_*i,i*_ = *λ*_*i*_, which are real and non-negative due to symmetry and positive semi-definiteness of **L**. Without loss of generality, we assume that the diagonal elements in Λ, and the corresponding columns in **U**, are sorted based on the magnitude of the eigenvalues, *i.e*., ∀*i, j* if *i < j* then Λ_*i,i*_ ≤ Λ_*j,j*_. As such, the graph Laplacian eigen-value set satisfies {0 = *λ*_1_ ≤ *λ*_2_ … *λ*_*N*_ := *λ*_max_ ≤ 2}, where the upper bound is guaranteed due to the use of the normalized Laplacian matrix (Chung, 1997). This set defines the Laplacian *spectrum* of the graph, and the eigenvector set {**u**_*i*_}_*i*=1,…,*N*_ defines an orthonormal basis that spans the ℝ^*N*^ space of vectors defined on the nodes of the graph; in the following, we occasionally refer to the Laplacian eigenvectors also as *eigenmodes*, a nomenclature commonly used in the neuroimaging community.

The eigenvalues of a graph Laplacian carry a notion of frequency, which is directly linked to the extent of spatial saliency manifested by their corresponding eigenvectors. To understand this link, a metric known as total variation (TV) (Sandryhaila and Moura, 2014) can be computed for each eigenvector, or more precisely, for any given graph signal. A graph signal defined on the nodes of a graph can be represented as a vector **x ∈ ℝ**^*N*^, where the *i*-th element, **x**[*i*], is the signal value at the *i*-th node of the graph. For a given graph signal **x**, the TV of **x** is defined as *TV* (**x**) = **x**^*T*^**Lx**, a measure that quantifies the extent of variation observed in **x** relative to the underlying graph structure. Given that the eigenvectors are orthonormal, *i.e*., 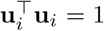 and that **Lu**_*i*_ = *λ*_*i*_**u**_*i*_, it follows that the TV of Laplacian eigenvectors reduces to 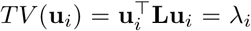, showing that the variability of each Laplacian eigenvector is reflected by the associated eigenvalue, or in other words, that Laplacian eigen-vectors associated to larger eigenvalues reflect a greater extent of spatial variability. Alternatively, spatial variability of graph signals/eigenmodes can be quantified via a measure of zero-crossings(Petrantonakis, 2021; Petrantonakis and Kompatsiaris, 2022). In particular, we define a weighted zero-crossing measure as (Abramian et al., 2021)

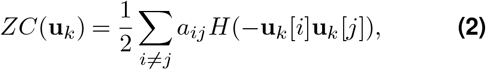

where *H*(·) is the Heaviside step function and *a*_*ij*_ is the edge weight that connects voxels *v*_*i*_ and *v*_*j*_. The higher the zero-crossing metric, the greater the associated variability in the eigenvectors’ spatial patterns.

### 2.C. Spectral decomposition of graph signals

For a given graph signal **x**, its spectral representation, commonly referred to as the graph Fourier transform (GFT) of **x**, is given as

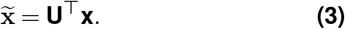

The signal can be perfectly recovered through the inverse GFT operation as, 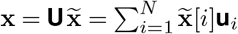. As such, any given graph signal can be seen as a linear combination of the orthonormal set of Laplacian eigenvectors. In particular, 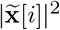 gives the energy spectral density of the signal associated to the *i*-th eigenmode. Given a set of graph signals 𝒳 = {**x**_*k*_ *}*_*k*=1,…,*S*_ (e.g. graph signals derived from individual time frames of a given fMRI session) we compute the ensemble energy spectral density (EESD) of the lower end of the spectrum of *𝒳* as

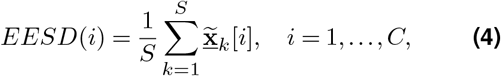

where *C* denotes a desired cutoff index specifying the number of lower end spectral indices to be studied (*C* = 1000 in this study), and 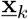 denotes the demeaned and normalized version of **x**_*k*_ obatined as

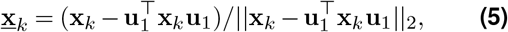

which ensures 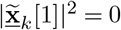 and 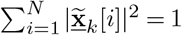.

### 2.D. Brain graph design

For each subject, we define a weighted brain graph characterized by a node set 𝒩= {1, …, *N*} defined based on the set of voxels that fall within the subject’s brain mask, covering gray matter (GM), white matter (WM) and cerebrospinal fluid (CSF), representing a 3D mesh arrangement. In particular, each node *i* is associated to a voxel, denoted *v*_*i*_, with coordinates (*x*_*i*_, *y*_*i*_, *z*_*i*_). The graph edges are defined based on the adjacency of voxels within the Moore neighborhood cubic lattice of size 3 × 3 × 3 and 5 × 5 × 5, where the latter size is only used in the ODF-based design. For the 5 × 5 × 5 design, voxels in the outer layer that fall in parallel to the voxels within the inner layer were excluded, enabling encoding of connections to a maximum of 98 voxels/directions in the neighborhood of each focal voxel, whereas the 3 × 3 × 3 design enables encoding connections to a maximum of 26 different directions. As such, the 5-connectivity design trades localization for better angular resolutions. With this definition of edges, each node *i* entails a neighborhood set of the indices of nodes in 𝒩 that are adjacent to it, denoted 𝒩_*i*_, where |𝒩_*i*_| ≤ 26 and ≤ 98 for the 3- and 5-connectivity designs, respectively.

We define the edge weights based on a measure of *inter-voxel fiber coherence* across all pairs of adjacent voxels. In particular, to make the presented method applicable to a wider range of available diffusion MRI data, we present two edge weighting schemes using two signal representation models, one using diffusion tensors and one using diffusion ODFs; we denote the resulting edge weighting schemes as the DTI-based and ODF-based methods, respectively. The tensor and ODF models both aim to represent structural information on the intra-voxel axon fiber arrangement. The diffusion tensor model can be seen as a multivariate Gaussian describing the distribution of fiber bundle alignment, whereas the diffusion ODF model defines the radial projection of the diffusion function, providing an estimate of the empirical distribution of water diffusion.

Let **r**_*ij*_ denote the vector pointing from the center of the voxel *v*_*i*_ to the center of the voxel *v*_*j*_. Let 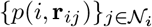 denote estimates of the extent of diffusion at voxel *v*_*i*_ in directions 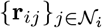. In the following, we first present a DTI-based and an ODF-based approach to estimate 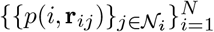. We then use these estimates to define the graph edge weights.

#### 2.D.1. DTI-based quantification of diffusion orientation

DTI is a model-based method for reconstructing the diffusion signal from diffusion MRI. The assumption in DTI is that the diffusion pattern follows the shape of a 3D ellipsoid. The molecular displacement of water at voxel *i* in the direction **r**_*ij*_ can be approximated by a 3D Gaussian distribution with real symmetric diffusion tensor **T** as the covariance matrix:

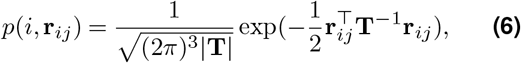

where |**T**| is the determinant of the diffusion tensor. The calculation of *p*(*i*, **r**_*ij*_) requires a discretization step that guarantees a one-to-one mapping between the (continuous) multivariate Gaussian model and the (discrete) weighting of nodes in the brain graph. For the 3 × 3 × 3 neighborhood encoding scheme, for a given voxel *v*_*i*_, the set of values 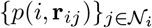 can be arranged into 3 × 3 × 3 discrete representation, which mimics the structure of a 3D finite impulse response (FIR) filter. As such, the problem of obtaining 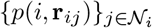 can be alternatively seen as that of obtaining the coefficients of an FIR filter. We provide the details of this procedure in the Supplementary Material.

#### 2.D.2. ODF-based quantification of diffusion orientation

Unlike diffusion tensors, ODFs do not follow a specific model and shape. Thus, a one-to-one discretization of a continuous function similar to that presented for the DTI model cannot be applied. Here we build on the construction previously presented by Iturria-Medina (Iturria-Medina et al., 2007). Within standard spherical co-ordinates, parametrized by (*r, θ, ϕ*,), let 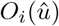 denote the ODF associated to voxel *v*_*i*_ with its center of coordinate being the voxel’s center, with 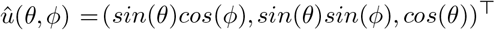 denoting the unit direction vector. Given 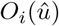, a measure of the extent of diffusion at voxel *v*_*i*_ along direction **r**_*ij*_ can be obtained as

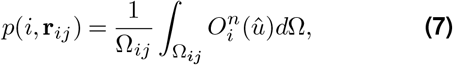

where Ω_*ij*_ denotes a given solid angle around **r**_*ij*_, *d*Ω = *sin*(*θ*)*dθdϕ* denotes the infinitesimal solid angle element; in particular, a solid angle of 4*π/*26 and 4*π/*98 is used for the 3- and 5-connectivity schemes, respectively. The exponent *n >* 0 is a desired power factor that is used to sharpen the ODF given the limited degree to which diffusion ODFs can differentiate fiber orientations (Jones et al., 2013). As such, *p*(*i*, **r**_*ij*_) gives an average measure of the surface area of the ODF within a spherical cap defined by the solid angle. Given a discrete representation of 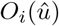 in form of *N*_*o*_ samples, denoted 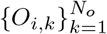, along *N*_*o*_ spherical directions from the center of voxels *v*_*i*_, (7) can be approximated as

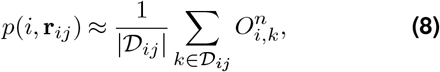

where 𝒟_*ij*_ denotes a subset of direction indices {1, …, *N*_*o*_} whose associated set of directions fall within Ω_*ij*_. The normalization by the cardinality of 𝒟_*ij*_ is due to the difference in the number of ODF samples that fall within the solid angle subtended along the different neighborhood directions.

#### 2.D.3 Brain graph edge weights

The graph edge weights *a*_*ij*_ are defined by using the estimates of diffusion orientation at the associated voxels *v*_*i*_ and *v*_*j*_, i.e., *p*(*i*, **r**_*ij*_) and *p*(*j*, **r**_*ji*_), as well as the strength of anisotropy at those voxels. In particular, for a given voxel *v*_*i*_, let *F* (*v*_*i*_) and *Q*(*v*_*i*_) denote the voxel’s fractional anisotropy (FA) and quantitative anisotropy (QA), respectively. FA can be calculated directly from the eigenvalues of the diffusion tensor (Alexander et al., 2007) and QA pertains to the amount of diffusion anisotropy along the fiber orientation as originally defined by Yeh et. al. (Yeh et al., 2010). Using these measures, we define the graph edge weights *a*_*ij*_ as

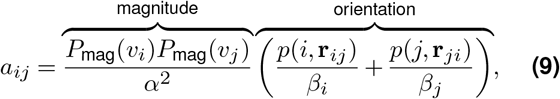

where 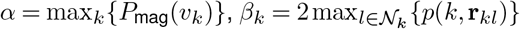 and *P*_mag_(*v*_*i*_) denotes the magnitude of anisotropy at voxel *i*, defined as

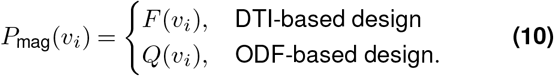

The connectivity structure of the graph is then characterized in **A**, such that *a*_*ij*_ *>* 0, given as in (9), if nodes *i* and *j* are connected through an edge, and *a*_*ij*_ = 0 if otherwise.

The *orientation* term in (9) gives a measure of diffusion orientation coherence between the two connected voxels, whereas the anisotropies give a *magnitude* measure that is useful in delineating tissues; the use of anisotropies is in contrast to using probabilistic tissue maps as used in (Iturria-Medina et al., 2007), enabling the design of the graphs using only the diffusion data. Given two adjacent voxels that exhibit highly coherent diffusion orientations, a large weight is associated to the connection only if the two voxels also exhibit notably large anisotropies. This interplay between the orientation term and magnitude term enables, for example, to prevent associating large weight to an edge between a WM voxel and a CSF voxel. Furthermore, the normalizations incorporated in the definition, i.e., the *α* and *β*_*k*_ terms, ensure having an unbiased definition of weights relative to the structure of the diffusion tensors/ODFs across the brain, and, mathematically, they impose bounds on the orientation and magnitude terms—both terms bounded to [0, 1], which in turn results in *a*_*ij*_ also being bounded to [0, 1].

### 2.E. Group-level eigenmodes

The voxel-wise nature of the studied graphs renders their size excessively large, with 7.8 ± 0.7 × 10^5^ nodes across the 100 subjects considered. The sheer size of the voxel-wise graphs impedes computing the full eigende-composition of the graph Laplacian, and, therefore, we compute and study the first leading 1000 eigenvectors corresponding to the lowest spectral frequencies. To preserve subtle subject-specific spatial details, we construct all graphs in the native space of each subject’s diffusion data. To enable inter-subject comparison of eigenmodes, the DARTEL normalization algorithm (Ashburner, 2007) implemented in SPM12 was used to define a group-level template coordinate space, based on the group’s T1-weighted MRI data. This results in a structural T1-weighted template as well as a set of transformation maps per subject. Each subject’s eigenmodes are then transformed into the template space using the subject-specific transformations, resulting in inter-subject spatially aligned eigenmodes.

### 2.F. Consistent inter-subject ordering of eigenmodes

The ordering of DARTEL-normalized eigenmodes is not necessarily consistent across subjects, including sign ambiguity and linear combinations between modes with close eigenvalues. To obtain a consistent ordering of the eigenmodes, and enable inter-subject comparison of individual eigenmodes, we used the Procrustes transform (Goodall, 1991), which finds the optimal rotation, translation, and/or reflection between two linear subspaces. We implemented a scheme of Procrustes transformation, where the subspace is defined by the first *K* DARTEL-normalized eigenmodes of any *M* subset of subjects. Let **u**_*i,m*_ denote the *i*-th eigenmode of subject *m*, and let *X*_*K,m*_ = [**u**_1,*m*_ **u**_2,*m*_ … **u**_*K,m*_].

#### Algorithm 1: group-level matching of eigen-modes

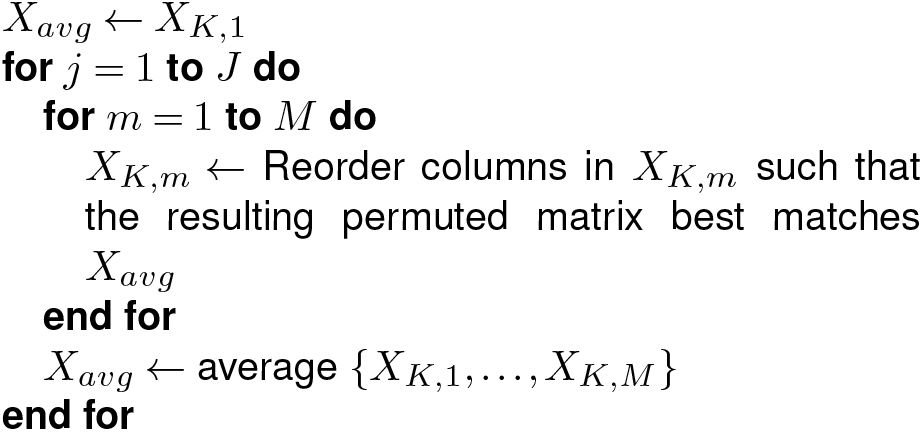

The *reordering* is done based on Procrustes transformation estimates, and *J* denotes the optimal number of iterations to ensure that *X*_*avg*_ does not remain biased towards its initial value, i.e., *X*_*K*,1_. The columns of the resulting 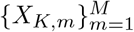 are reordered in such a way that they optimally match each other and can thus be compared across the subjects. Although the Procrustes transformation removes much of the variance that is common between subjects, it cannot discount for subject-specific details; in the following section we define a metric that quantifies the extent of remaining inter-subject structural variability.

### 2.G. Quantification of the extent of inter-subject structural variability as encoded in brain graphs

The precision of the Procrustes transformation can be evaluated by quantifying the cosine similarity between corresponding eigenmodes of different subjects, after DARTEL normalization and Procrustes transformation. For any pair of subjects, a symmetric cosine similarity matrix is obtained, where the deviation of the off-diagonal elements of the matrix from zero quantifies inter-subject structural variability. If the set of eigenmodes of two subjects has been ideally matched, the cosine similarity matrix should be the identity matrix. Therefore, to quantify the mismatch between a pair of subjects *m* and *n* based on their first *K* eigenmodes, we define a measure of the extent of inter-subject structural variability, termed *Procrustes error*, via computing the cosine similarity between pairs of eigenmodes as

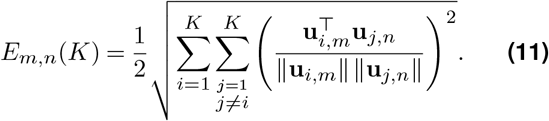

We used this error term to determine *J* in Algorithm 1 and also to compare the different graph designs.

To validate the performance of the Procrustes transformation, we implemented a bootstrap scheme, which successively applies the transformation on the first K eigen-modes of two randomly chosen subjects; the implementation is summarized in the following algorithm:

#### Algorithm 2: Procrustes validation

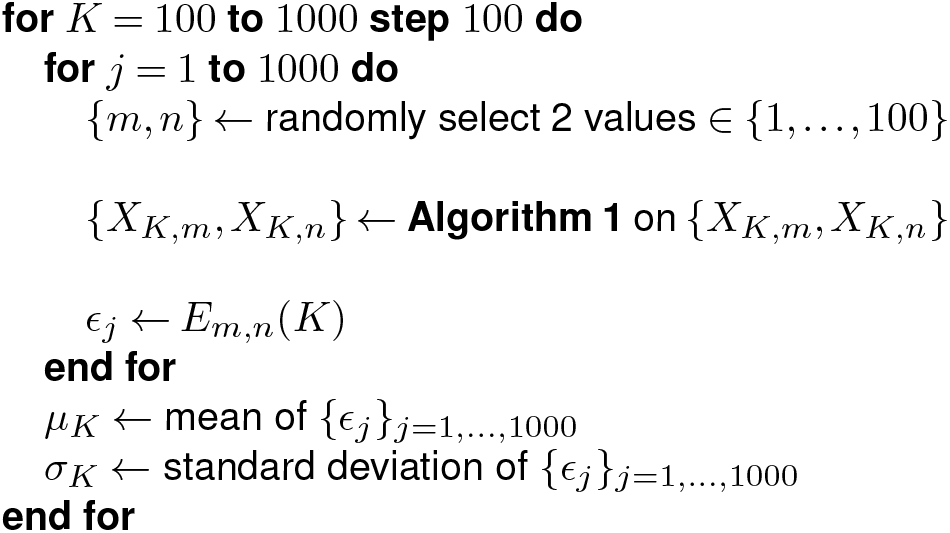

When comparing two brain graph designs, the design that results in a higher *µ*_*K*_, with reasonably small *σ*_*K*_, is interpreted as capturing more subject-specific structural features.

### 2.H. Spectral decomposition of fMRI data

As proof-of-concept of the applicability of the proposed whole-brain, voxel-wise brain graphs, we evaluated the extent to which brain fMRI data are spatially shaped by the underlying brain structure as encoded by the graphs. In particular, we constructed fMRI graph signals from six functional tasks as well a resting-state acquisition, across 100 subjects. Each fMRI time frame was transformed in to a single graph signal. This was done by extracting the fMRI voxels associated to the graph vertices, i.e., voxels that fall within each subject’s brain mask, arranging them as a vector, ordered based on the order of vertices as reflected in each subject’s graph adjacency matrix. As such, for each subject, a time-evolving series of graph signals were obtained from each of the subjects’ task or resting-state 4D fMRI volumes. We studied the energy spectral density of the extracted graph signals associated to the first 1000 spectral indices of the ODF-3 graph. To serve as a null, we also evaluated the energy spectral density of synthesized shuffled fMRI signals— obtained through random permutation of voxel indices of each fMRI volume to destroy spatial order, as well as white Gaussian noise signals.

## 3. Results

Figure 1 shows the degree distributions of the graphs across the different tissue types. The connectivity strength is highest in WM nodes compared to GM and CSF nodes. The degree distribution of ODF-5 graphs is a shifted version of that of the ODF-3 graphs towards a higher degree, reflecting the larger number of connections possible with the ODF-5 design. It can be observed that the nodal degrees within gray matter and CSF almost coincide for the DTI design whereas this is much less pronounced in the ODF designs (compare Figure 1(A) with Figs. 1(B) and (C)); this is more pronounced for the ODF-3 design, which reflects that the ODF-3 design bears larger differences across tissue types compared to the DTI-3 design. Figure 1(D) shows the distribution of nodes for the different tissue types across all subjects; the median number of nodes for GM, WM, and CSF were 254 299, 221 964, and 294 028, respectively, with standard deviations 25 322, 28 579, and 25 742, respectively.

**Figure 1.**
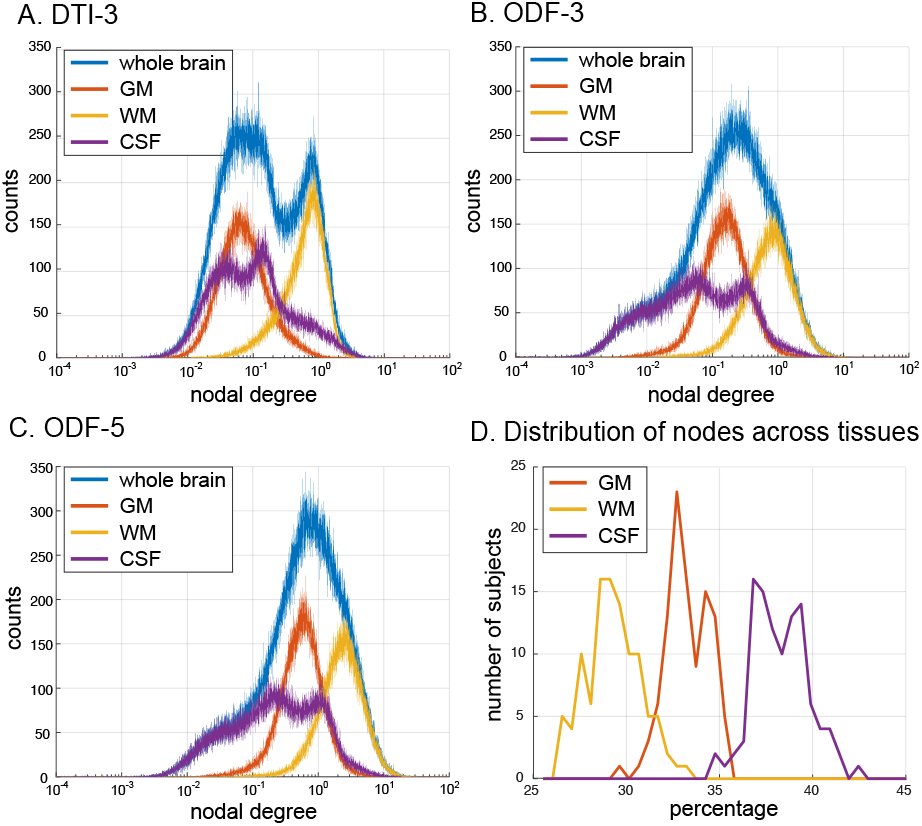
Degree distribution of the (A) DTI-3, (B) ODF-3, and (C) ODF-5 voxel-wise brain graphs for a representative subject. (D) Distribution of the number of nodes in each tissue type across all subjects.

### 3.A. Spectral Comparison

Figure 2(A) shows the first Laplacian eigenmodes obtained from the three brain graphs of a representative subject. Noting that the first eigenmode of **L** is a function of the graph nodal degrees—**u**_1_ = **D**^1*/*2^**1** where **1** denotes the constant function that assumes the value of 1 on each node, the spatial pattern manifested by the first eigenmodes is a corroboration of the results shown in Figure 1(A)-(C), demonstrating that the distinction between tissue types naturally arises from the assignment of the connectivity weights in the brain graph, in particular through the assignment of the magnitude term in (9). Moreover, the first eigenmode manifests a specific profile of local tissue structure, in which higher values reflect voxels/regions that are more strongly connected to their surrounding neighbourhood, particularly observed at regions of less ambiguous fiber structure, e.g. within the corpus callosum The second and third eigenmodes shown in Figure 2(B) manifest global morphological organization of the brain, contrasting the posterior and anterior, and the left and right brain regions, respectively. The next eigenmodes exhibit a greater extent of spatial variability and localized information.

**Figure 2.**
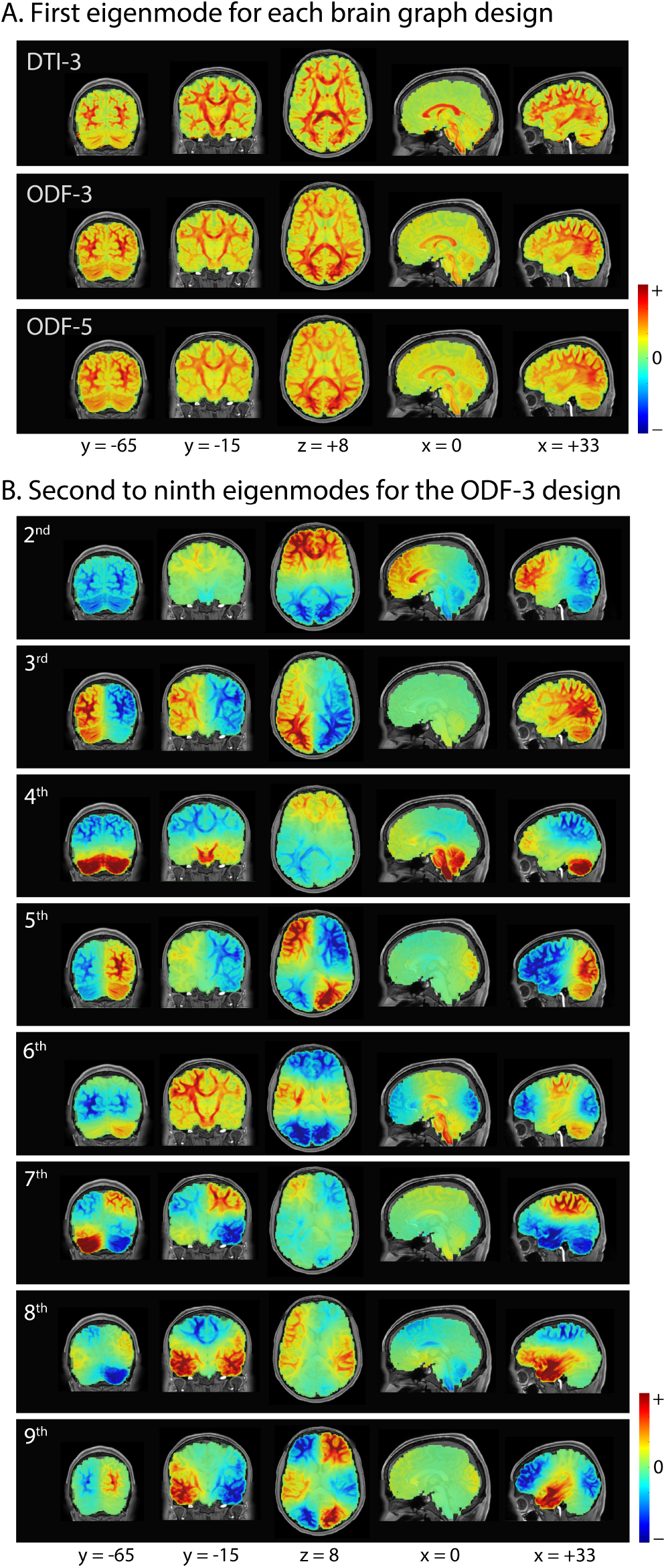
(A) First Laplacian eigenmode of 3-connectivity DTI, 3-connectivity ODF and 5-connectivity ODF brain graphs of a representative subject. (B) The next lowest frequency eigenmodes corresponding to the 3-connectivity ODF brain graph.

Figure 3(A) shows the lower-end graph spectra in which the rate of increase in the first 1000 eigenvalues are shown. The ODF-5 graph has relatively larger eigen-values than the ODF-3 and DTI-3 graphs. Given that the eigenvalues entail a notion of spatial saliency, the increase in local connectivity in the ODF-5 implies that the associated eigenmodes have greater degrees of freedom, and as such, spatial saliency can become higher.

**Figure 3.**
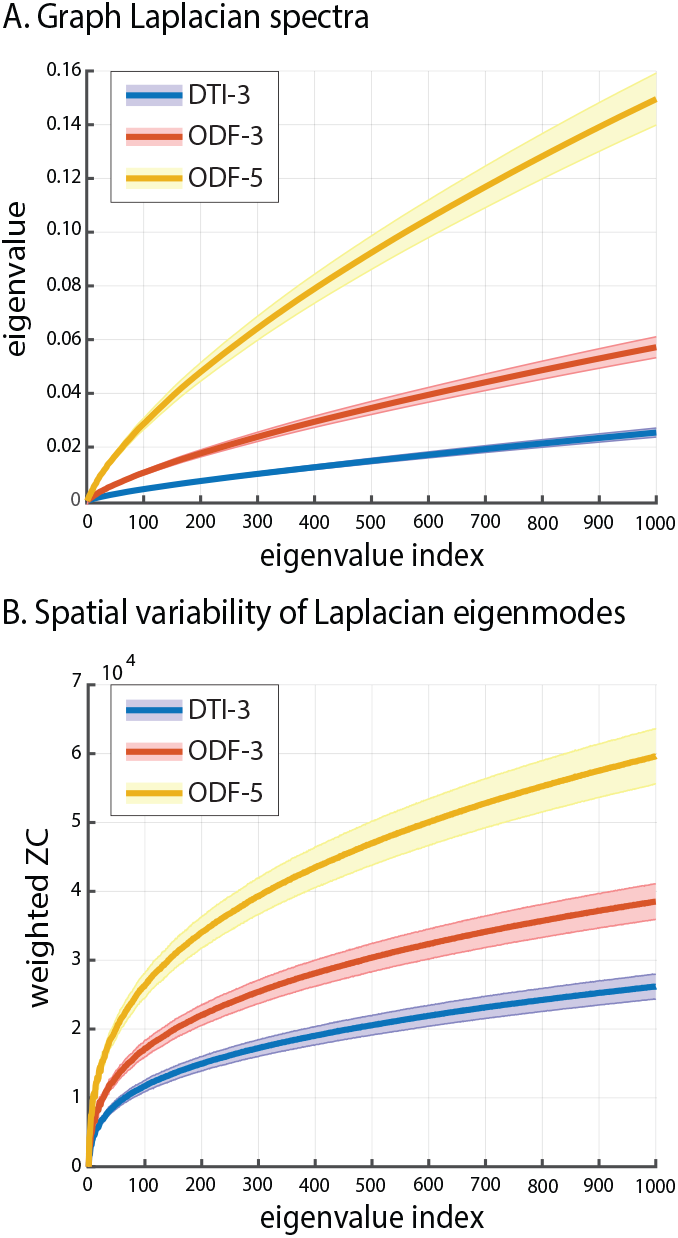
(A) Lower-end eigenvalues of DTI-3, ODF-3 and ODF-5 graphs, consisting of their first 1000 eigenvalues. (B) weighted zero-crossing the corresponding eigenmodes; cf. (2); solid lines show the mean and shades show the standard deviation across the 100 subjects.

The spatial saliency of the eigenmodes’ can be quantified by computing their weighted zero-crossing, cf. (2). Figure 3(B) shows a trend that is consistent with that of the eigenvalues, thereby confirming the general notion that higher indexed eigenmodes encompass a larger extent of spatial saliency. Graph Laplacian spectra

### 3.B. Procrustes Validation

We evaluated the inherent inter-subject variability encoded in two voxel-wise brain graph designs via the Procrustes transform, which finds the optimal configuration that matches single subject eigenmodes to an averaged set. This step, however, cannot account for subject-specific fiber pathways encoded in the eigen-modes, as manifested by the cosine similarity analysis of pairs of subjects. The cosine similarity matrices of two sets of eigenmodes before and after applying Procrustes transformation are shown in Figure 4. The diagonal structure of the cosine similarity matrix associated to the transformed eigenmodes shows the effectiveness of the transformation in aligning eigenmodes of the same neuroanatomical spatial nature. The insets show traces of flipped signs and unordered eigenmodes in the original vectors, which were corrected after Procrustes transformation. The precision of the Procrustes analysis improves after several rounds of the transformation; see Figure S1 in Supplementary Materials.

**Figure 4.**
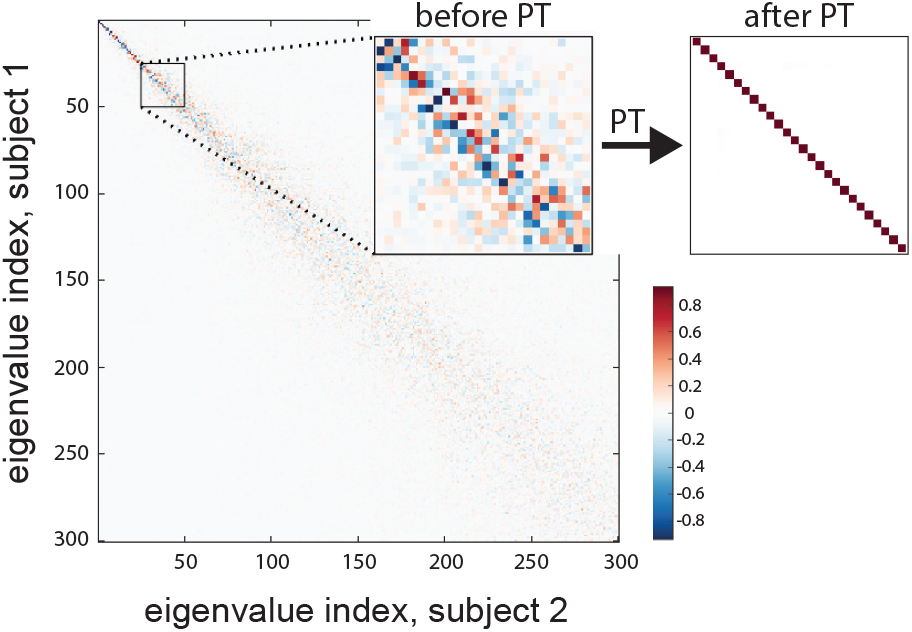
Cosine similarity between the first 300 eigenmodes before and after Procrustes transformation (PT) for two representative subjects.

Figure 5 shows the Procrustes error when different subset of eigenmodes are used, for each of the three graph designs, reflecting the extent of inter-subject structural variability captured by the eigenmodes. Furthermore, to serve as a null for comparing and validating the brain graphs, we synthesized 100 random orthonormal vectors of the same dimension as each of the subject-specific eigenmodes, applied DARTEL normalization, and then subjected the resulting vectors to Procrustes validation. All three brain graphs show a decreasing trend, and they get closer to that of the null upon reaching higher *K*-values. Despite the differences in the Procrustes errors across the three brain graphs for low values of *K*, all errors eventually converge to a single point for high *K*. In addition, the ODF-based designs show higher Procrustes errors than the DTI-based design, across K. The ODF-5 design slightly outperformed the ODF-3 design, suggesting the benefit of using the larger neighborhood in encoding fiber orientations with better angular resolution.

**Figure 5.**
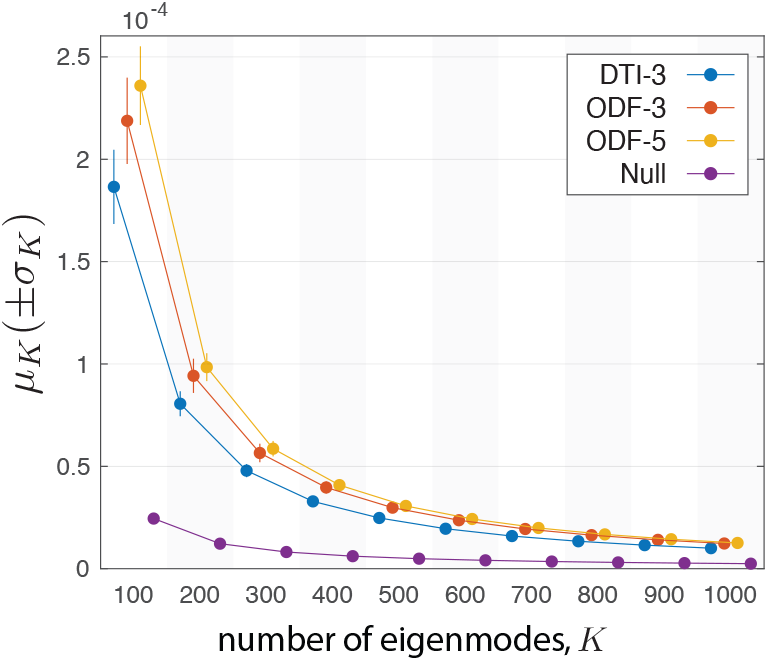
Quantification of the extent of inter-subject structural variability captured by the eigenmodes, cf. Algorithm 2. The four markers (DTI-3, ODF-3, ODF-5, and Null) within each vertical shade are associated to the same K.

### 3.C. Spectral decomposition of fMRI

Figure 6(A) shows the ensemble energy spectral densities of the fMRI. The energy spectral density of the fMRI data is characterized by a power-law behavior. The lowest frequency component eigenmodes capture the majority of the energy content (spatial variability), which is about two orders of magnitude greater than what is captured by the 1000th eigenmode. Moreover, a notable dispersion is observed in the energy profiles of approximately the first 100 spectral indices across the different tasks, whereas for the higher spectral indices, the profiles are more closely packed, following a steady power-law drop. This observation is more apparent by inspecting the cumulative ensemble energy (CEE) profiles, see Figure 6(B); results are shown also for synthesized shuffled fMRI signals (cf. Section H) and Gaussian noise signals. In particular, the CEE profiles of task and rest fMRI sharply differ from that of Gaussian noise as well as shuffled fMRI data. For Gaussian noise, each eigenmode captures a fraction of the total energy equivalent to approximately 1*/N*, where *N* is the number of the graph nodes, whereas shuffled fMRI still entails the distribution of values as in the real fMRI data, but lacks the exquisite spatial dependencies manifested in fMRI data that is linked to the underlying structure. The CEE profiles of fMRI data show that functional brain activity is expressed preferentially by lower-frequency components; approximately 85% of the total signal energy content^4^ captured by the first 1000 eigenvectors, wherein the contributions from the first eigenmode are equal to zero due to that the data were demeaned, cf. Section **??**.

**Figure 6.**
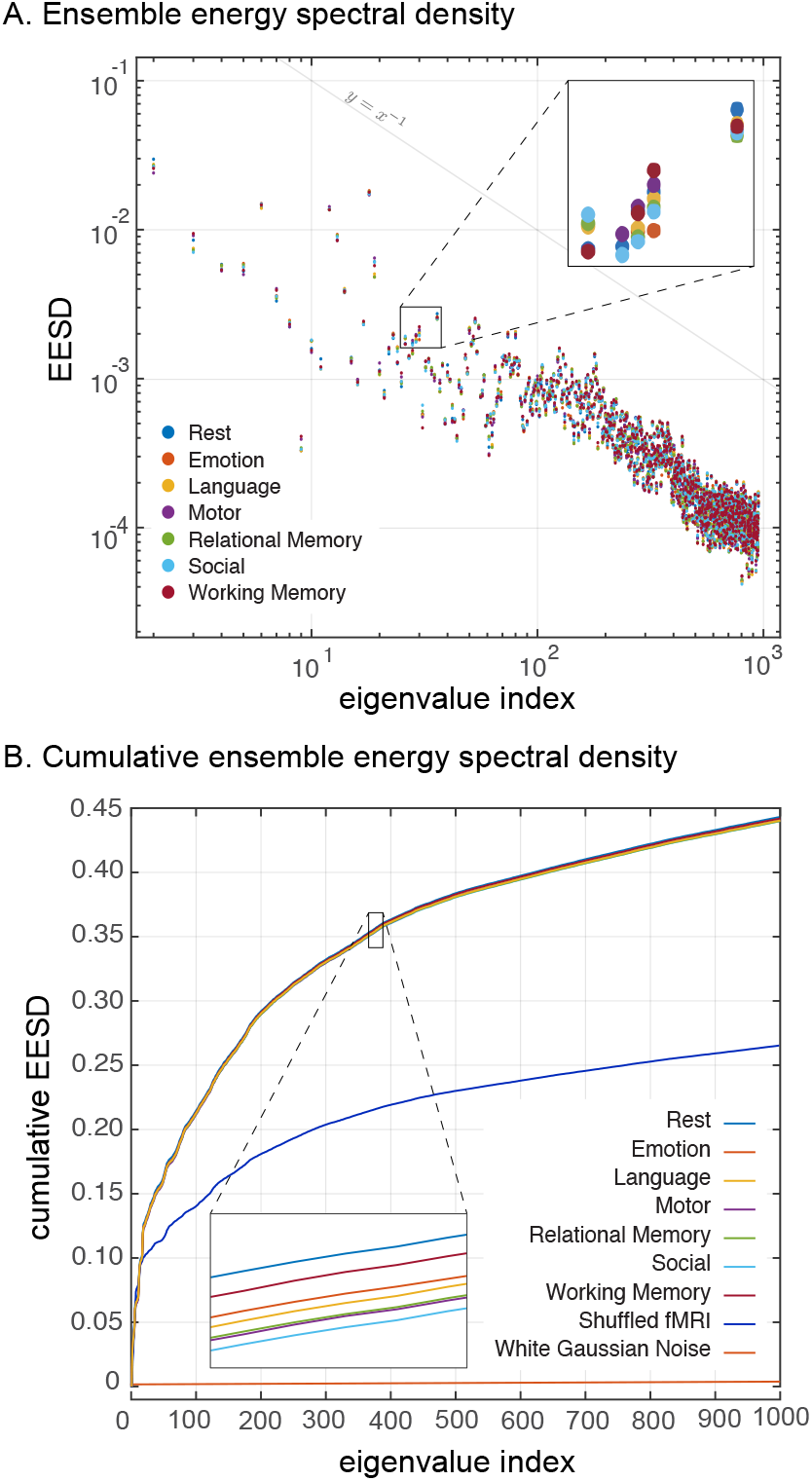
(A) Ensemble average energy spectral density of fMRI data of 100 subjects for the first 1000 eigenmodes of ODF-3 brain graph. (B) Same as in (A) but showing the cumulative values as well as a comparison to shuffled fMRI data and white Gaussian noise.

## 4. Discussion

Voxel-wise brain graphs enable overcoming several limitations associated to existing region-wise brain graphs. For example, they obviate the need for cortical parcellation, and as such, downstream analysis will not be affected by the choice of parcellation scheme (de Reus and Van den Heuvel, 2013). Moreover, analyses on local-encoding voxel-wise graphs prevents variations in results as a function of the algorithm that is employed to approximate the number of WM tracts for region-wise graphs (Maier-Hein et al.). Lastly, given that the proposed graphs are constructed at the native diffusion space and encompass the whole brain, tissue segmentation and subsequent transformation to a template space is not needed, which can be challenging especially in populations that exhibit a complex mixture of brain structural deficits.

Results on fMRI show that despite the high dimensionality of voxel-wise graphs, brain functional activity can be well approximated by merely a small subset of their low-frequency Laplacian harmonics, whereas in contrast for region-wise brain graphs a larger subset of their total number of Laplacian harmonics is required (Atasoy et al., 2017; Preti and Van De Ville, 2019; Glomb et al., 2020). This observation shows that functional brain maps are smooth relative to the underlying fiber architecture and tissue profile morphologies, which, on the one hand, is consistent with energy profile of fMRI graph signals on tissue-specific, voxel-wise graphs (Behjat et al., 2015, 2021), and on the other hand, can be linked to the decreasing trend observed in the Procrustes validation errors (see Figure 5), where increasing K-values reduce the error close to that of a randomly generated graph. This energy pattern is reminiscent of a power-law behaviour, which is interesting in light of the evidence of scale-free behaviour in human brain activity (Van De Ville et al., 2010; Ciuciu et al., 2012) observed in both temporal and spatial scales. Moreover, a practical implication of such energy profiles is that the lower-spectral-end energy content of fMRI data on whole-brain voxel-wise graphs has the potential to provide signatures of mental activity, similar to that observed for cerebral cortex gray matter graphs (Behjat and Larsson, 2020), which substantially reduces the computational burden associated to diagonalization of the graph Laplacian. Alternatively, to study the energy profile across the spectrum, a filter design scheme that adapts to the ensemble signal content can be used to efficiently partition the spectrum (Behjat et al., 2016), which can be implemented in a computationally efficient manner (Behjat and Van De Ville, 2019), obviating the need to even compute individual eigenvectors.

Voxel-wise brain graphs hold the potential to open new research avenues to study the brain. One avenue is to study the graphs from a pure structural perspective, using spectral graph theoretical measures that have been used to e.g. discriminate auditory gyri subtypes (Maghsadhagh et al., 2021), or to perform subject identification and characterization of hemispheric asymmetries (Wachinger et al., 2016). A second, more interesting, avenue of research is to employ voxel-wise brain graphs within the context of relating brain structure to function. The proposed voxel-wise graphs can be leveraged to perform whole-brain anatomically-informed spatial filtering and interpolation of fMRI data, operations that are inherent within numerous fMRI processing pipelines; e.g. spatial smoothing to enhance whole-brain fMRI activation mapping, as done using tissue-specific designs in gray matter (Behjat et al., 2015, 2021) and white matter (Abramian et al., 2021; Zhao et al., 2022). Moreover, it yet remains to be studied how functional connectivity (FC) and their associated measures can be extended to accurately integrate structural information. FC has often been associated to Euclidean distance (Alexander-Bloch et al., 2013), whereas by using the Laplacian eigen-modes of the proposed graph, functional distance can be better interpreted in relation to the underlying brain structure. That is, functional variations that are captured using low-frequency eigenmodes are smooth with respect to long-distance white matter bundles, whereas localized and short-distance functional associations are expected to be dominated by higher frequency components. Lastly, the exquisite voxel-wise scale of the proposed graphs can enable assessing the extent to which brain structural-functional relations hold at spatially finer mesoscales (Mansour-L et al., 2021); e.g. by using graph Slepians (Van De Ville et al., 2017; Petrovic et al., 2019; Georgiadis et al., 2021) or variants of localized graph filter banks (Shuman, 2020; Isufi et al., 2022) and spectral transforms (Ghandehari et al., 2021; de Loynes et al., 2022; Tay, 2022), focus can be placed on a particular subset of nodes, thus, providing a finer level of analytical resolution than that provided by conventional region-wise graphs.

## 5. Conclusion

Two methods for constructing voxel-wise brain graphs from diffusion MRI data were studied. Through a Procrustes validation scheme that reflects inter-subject structural differences, it was shown that low-frequency eigenmodes of such high spatial resolution graphs reflect the highest amount of structural information from diffusion MRI. This finding was corroborated by the manifested energy spectral density of functional signals showing the preferential expression of human brain activity onto lower frequency components. Overall, the presented results signify the capability of voxel-wise brain graphs’ eigenmodes in capturing anatomically-constrained functional variations that are specific to different cognitive tasks. By treating voxel-wise brain graphs as the scaffold on which brain function is observed, they hold the potential to open new research avenues to study the brain, in particular, enabling the development of novel GSP methods to study the interplay between brain structure and function, in health and disease.

## Acknowledgements

This work was supported in part by the Swiss National Science Foundation under Grant 205321-163376 and in part by the Swedish Research Council under Grant 2018-06689. A preliminary version of this work was presented in (Tarun et al., 2019).

## Supplementary Materials

### Iterative Procrustes analysis

The precision of the Procrustes analysis improves after several rounds of the transformation. The evolution of the Procrustes error for each successive iterations of Procrustes transformation, evaluated through the term *µ*_*K*_ introduced in Algorithm 2, is displayed in Fig. S1(A). After the first iteration of Procrustes transformation, a large is observed in the error (about 60% change); after three iterations (for *J* = 3, 94% drop of error), we obtained the final set of group-averaged eigenmodes, the first few of which are displayed in Fig. S1(B). Major WM features that are relatively smoother than subject-specific bundles are preserved, whereas subject-specific local structures are lost.

**Figure S1.**
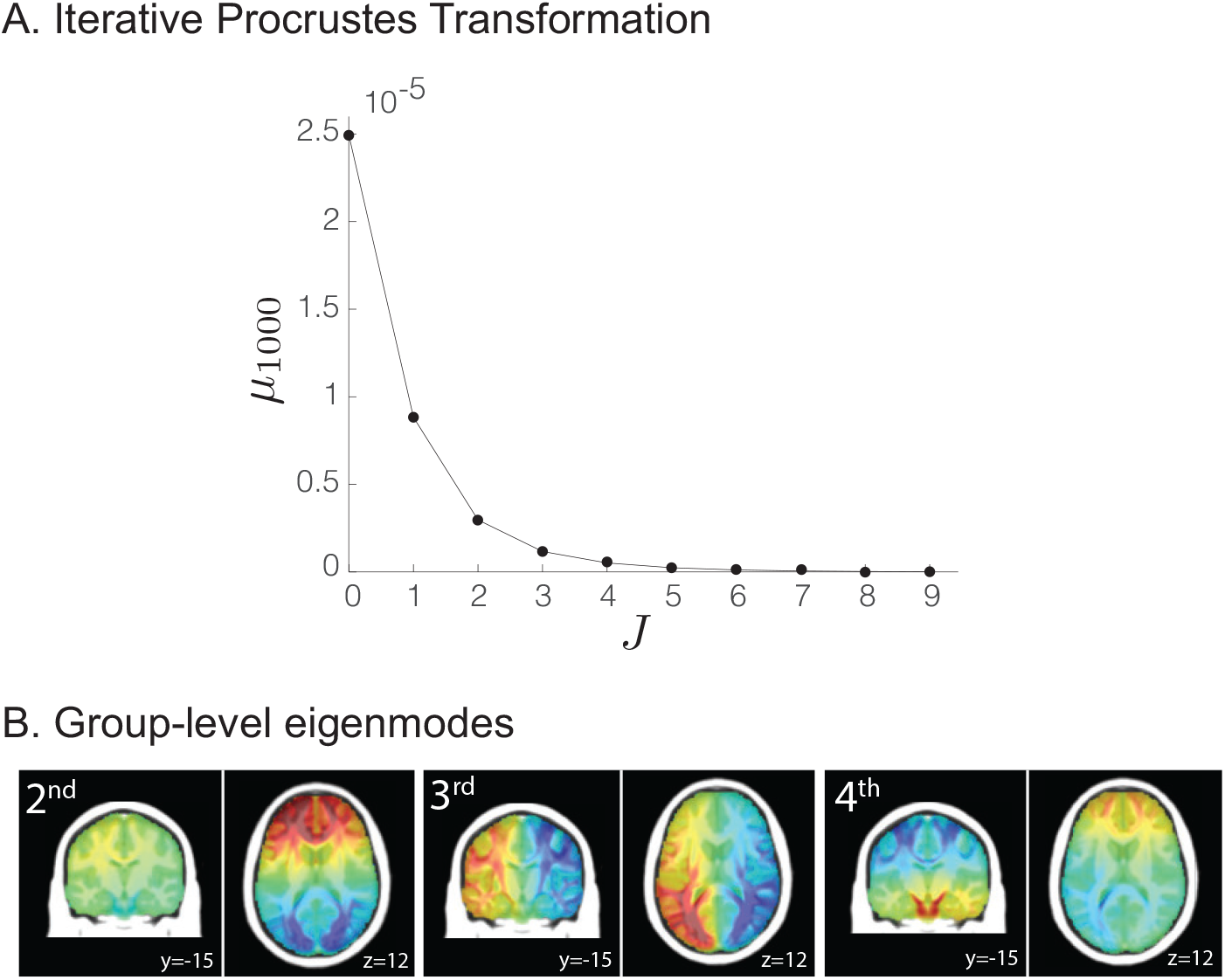
(A) Residual error after successive PTs, cf. Algorithm 1. (B) The second, third, and fourth group-averaged eigenmode.

### Cartesian discretisation of a Gaussian via FIR filter design

The molecular displacement of water at voxel *i* in the direction **r**_*ij*_ can be approximated by a 3D Gaussian distribution with real symmetric diffusion tensor **T** as the covariance matrix:

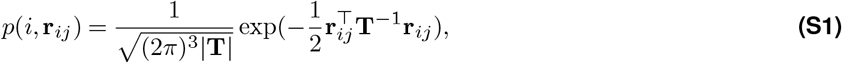

where |**T**| is the determinant of the diffusion tensor. The calculation of *p*(*i*, **r**_*ij*_) requires a discretization step that guarantees a one-to-one mapping between the (continuous) multivariate Gaussian model and the (discrete) weighting of nodes in the brain graph. Note that the Fourier transform of the Gaussian function in (S1) is another Gaussian function. For the 3 *×* 3 *×* 3 neighborhood encoding scheme, for a given voxel *v*_*i*_, the set of values 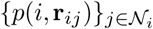 can be arranged into 3 *×* 3 *×* 3 discrete representation, which mimics the structure of a 3D finite impulse response (FIR) filter. Let **h**[*·, ·, ·*] denote an FIR filter, characterized by it’s frequency response *H*(*ω*) as

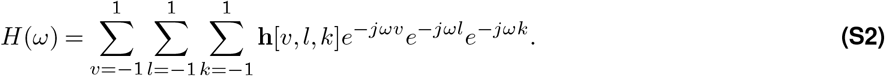

The coefficients **h** can then be obtained by a one-to-one mapping from the continuous domain to the discrete domain through the matching of corresponding frequencies. The frequency matching is done by considering the cartesian discretization of a Gaussian using its Fourier domain representation. For simplicity of notation, we present the discretization process for a 1D Gaussian, which can be readily extended to higher dimensions. The 1D Gaussian is given as

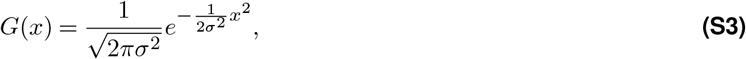

where *σ*^2^ is a constant which describes the variance of the curve and 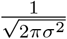 is a normalization factor. Its Fourier transform is given by

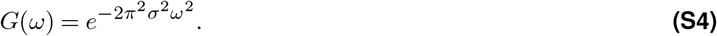

Obtaining a 3-sample discretization of the Gaussian using it’s Fourier domain representation is a problem that can be equivalently seen as finding the coefficients of a 3-tap FIR filter, **h**, matching the Frequency response given in *G*(*ω*) with that of the filter ℋ(*ω*) given as

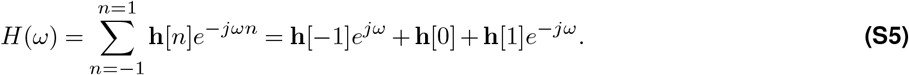

Noting that 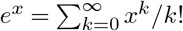, Taylor series expansion of *G*(*ω*) and *H*(*ω*) up to the second order gives

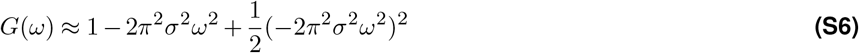

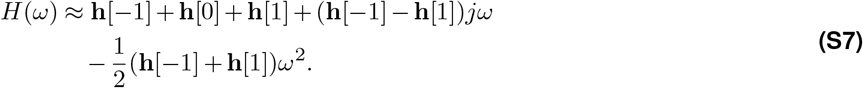

Matching the coefficients of the polynomials in (S6) and (S7), a system of linear equations is obtained as

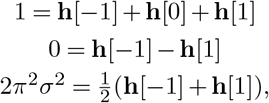

Where the solution gives the desired FIR impulse response as

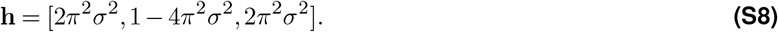

In the case of the 3D multivariate Gaussian density, we have 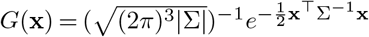, wherein the exponent is a quadratic form in the 3D vector variable **x** covering *x*–, *y*–, and *z*–directions, and *σ*^2^ is replaced by a 3 × 3 invertible covariance matrix Σ that is equivalent to the tensor **T** in (S1). The goal is then to find the coefficients of a 3 × 3 × 3 FIR filter by matching its frequency response ℋ(*ω*), as given in (S2), to the Fourier transform of the 3D Gaussian, which can be derived in a similar way as done for the 1D case above.

## Footnotes

1 https://www.fil.ion.ucl.ac.uk/spm/software/spm12

2 https://fsl.fmrib.ox.ac.uk

3 https://dsi-studio.labsolver.org

4 For each graph signal, the first eigenmode captured approximately 70% of the total signal energy. In the demeaning step, cf. Section **??**, the contribution from the first eigenmode was regressed out. As such, baed on Figure 6(B), the total amount of energy captured by the first 1000 eigenmodes amounts to approximately 85% of total signal ener-gies of the original signals, i.e., 70 + 0.45 × 30.

